# A red fluorescent protein with improved monomericity enables ratiometric voltage imaging with ASAP3

**DOI:** 10.1101/2020.10.09.328922

**Authors:** Benjamin B. Kim, Haodi Wu, Yukun A. Hao, Michael Pan, Mariya Chavarha, Michael Westberg, Francois St-Pierre, Joseph C. Wu, Michael Z. Lin

## Abstract

A ratiometric genetically encoded voltage indicator (GEVI) would be desirable for tracking transmembrane voltage changes in cells that are undergoing motion. To create a high-performance ratiometric GEVI, we explored the possibility of adding a voltage-independent red fluorophore to ASAP3, a high-gain green fluorescent GEVI. We performed combinatorial multi-site mutagenesis on the cyan-excitable red fluorescent protein mCyRFP1 to enhance brightness and monomericity, creating mCyRFP3. Among red fluorescent proteins tested, mCyRFP3 proved to be the least perturbing when fused to ASAP3. We demonstrate that the red fluorescence of ASAP3-mCyRFP3 (ASAP3-R3) provides an effective reference channel to remove motion artifacts from voltage-induced changes in green fluorescence. Finally we use ASAP3-R3 to visualize membrane voltage changes throughout the cell cycle of motile cells.

## Introduction

Ratiometric genetically encoded voltage indicators (GEVIs) are needed to remove unwanted artifacts of membrane motion so that changes in membrane potential can be discriminated from motion-induced changes in fluorescence. Membrane motion occurs when cells migrate or change shape, for example when cultured cardiomyocytes contract. Membrane motion is also an inherent feature of contractile tissues such as skeletal muscles, the gastrointestinal tract, or the heart, and sample movement relative to the camera is an omnipresent artifact during microscopy in live animals due to breathing, hemodynamic motions, or animal movement^1^.

A properly designed ratiometric GEVI would be able to provide a readout corrected for changes in indicator abundance caused by motion if two criteria are fulfilled. First, a reference fluorophore’s brightness must be either independent or inversely correlated to that of a voltage-modulated fluorophore of a different wavelengths in the same molecule, so that the ratio of emissions at the two wavelengths at any point in space can be related to voltage independently of indicator abundance at that point. Second, both the fluorophores should be excitable by the same wavelength of light, as this would enable simultaneous recording of the two fluorophores in separate channels, thereby increasing speed, simplicity, and precision. GEVIs that utilize a change in FRET efficiency between two fluorescent proteins meet these requirements, but engineering large responses in FRET-based GEVIs has proven to be difficult^2^.

Another solution may be to use a voltage-independent fluorophore with a large Stokes shift that can be excited at the same wavelength as the voltage sensitive-fluorophore while keeping their emissions easily separable. Many recently developed GEVIs with high voltage responsivity function via brightness modulation of a single fluorophore. For example, ASAP-family GEVIs utilize a single circularly permuted green fluorescent protein (GFP) domain^3–6^. We sought to create a high-performance ratiometric GEVI by fusing a voltage-independent large-Stokes-shift (LSS) red fluorescent protein (RFP) to ASAP3.

While many LSS RFPs have been developed, they suffer from residual oligomerization^7,8^ or poor brightness^9^. Here, we report the engineering of mCyRFP3, a brighter and more monomeric variant of the mCyRFP3 cyan-excitable RFP^8^. We show that mCyRFP3’s photophysical properties are similar to those of the parent dimer CyRFP1 while demonstrating improved brightness and maturation in mammalian cells. We demonstrate that mCyRFP3 is unique in allowing single-excitation dual emission imaging with efficient membrane localization when fused to the green fluorescent ASAP3. We further demonstrate that the green/red ratio of ASAP3-mCyRFP3 (ASAP3-R3) reports voltage changes in beating cardiomyocytes more reliably than ASAP3 alone. Finally, we use ASAP3-R3 for optical recording of membrane voltage changes during the cell cycle.

## Results

To create a ratiometric GEVI, we initially fused ASAP-family GEVIs at their C-terminus to the cyan-excitable red fluorescent protein CyRFP1 or its more monomeric variant mCyRFP1. The cyan excitation of CyRFP1 and mCyRFP1 allows them to be co-excited efficiently with ASAP1 by 488-nm light (or 950-nm light in two-photon mode). We also tested mRuby3, FusionRed, and mCherry, which have been fused to a variety of proteins without causing toxicity^10,11^. Disappointingly, all five fusions accumulated in the cell body and dendrites in structures too large to represent single vesicles, suggesting protein aggregation (**Fig. 1**). These accumulations were more visible in the red channel than the green channel, likely because the RFPs are located in the pH-neutral cytosol while the GFP moiety of ASAP is located in the acidic lumen of the secretory pathway. The resulting differences in green/red ratios across the cell prevents the use of the red channel to correct for movement artifacts. In addition, GEVI molecules trapped in intracellular accumulations will not respond to voltage changes at the membrane. As the CyRFP1 fusion appeared to be better localized to the membrane compared to the other RFP fusions, we chose CyRFP1 as a template for creating a more monomeric RFP.

**Figure 1.**
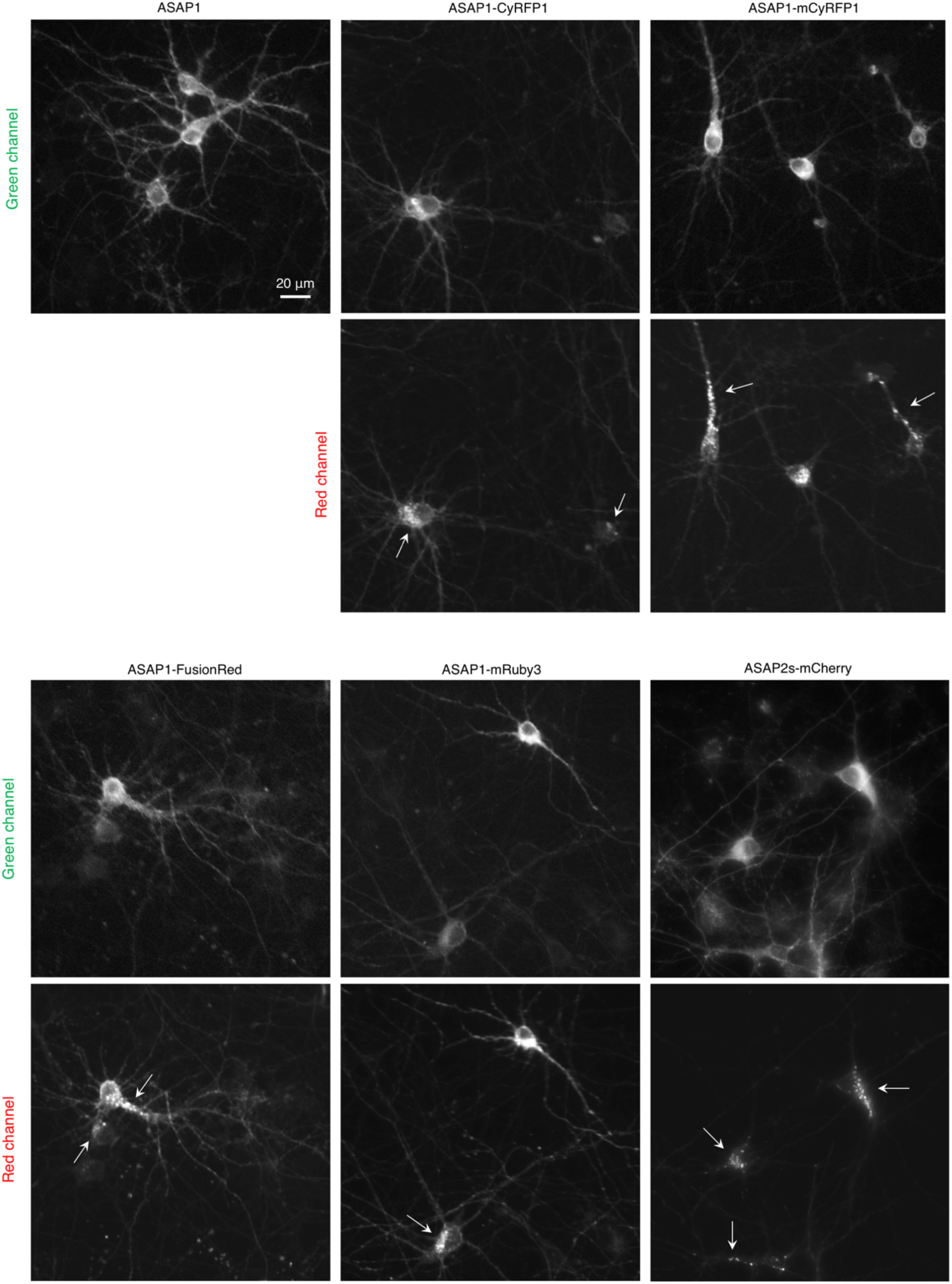
Initial candidates for ratiometric ASAP expressed in rat hippocampal neurons. Arrows point to large accumulations of the RFP-tagged ASAPs in the cell body or neurites.

mCyRFP1 had shown increased monomericity relative to CyRFP1 *in vitro* due to a surface A161K mutation, which breaks a hydrophobic contact with Tyr-148 across the cross-dimer interface^8^ **(Fig. 2a)**. We hypothesized that A161K may not be the optimal mutation for breaking this hydrophobic contact, e.g. if retaining tyrosine at position 148 is not ideal for folding or hydration in the monomeric state. Indeed, performing double saturation mutagenesis and screening for brightness yielded a Y148T A161V mutant that was more monomeric in vitro (**Fig. 2b**). We designated this protein mCyRFP3, since the name mCyRFP3 was already assigned to a different mCyRFP1 variant^12^. mCyRFP3 exhibited absorbance and excitation centered near 500 nm and peak emission of 586 nm, slightly red-shifted from mCyRFP1 **(Fig. 2c)**. mCyRFP3 showed improved fluorescence quantum yield of 0.72 and an extinction coefficient of 38 mM-1 cm-1 **(Table 1)**, making its molar brightness more than 50% brighter than its predecessor mCyRFP1. mCyRFP3 fluorescence demonstrated a pKa of 4.1 and was extremely stable in the physiological pH range **(Fig. 2d)**. mCyRFP3 photobleached with a normalized half-time of 75 s **(Fig. 2e)**, and exhibited a maturation half-time of 12.5 min **(Fig. 2f)**.

**Table 1.** Characteristics of mCyRFP3.

|  | Peak<br>excitation<br>(nm)<sup>a</sup> | Peak EC<br>(mM<sup>-1</sup>cm<sup>-1</sup>) | Peak<br>emission<br>(nm) | QY | Brightness<br>(mM<sup>-1</sup>cm<sup>-1</sup>) | pK<sub>a</sub> | Photostability<br>(s) |
| --- | --- | --- | --- | --- | --- | --- | --- |
| CyRFP1 | 528 (488) | 48 | 591 | 0.75 | 36 | 6.1 | ND |
| mCyRFP1 | 528 (488) | 27 | 594 | 0.65 | 17 | 5.8 | 40 |
| mCyRFP3 | 522 (488) | 38 | 588 | 0.72 | 28 | 4.8 | 70 |

<sup>a</sup>A minor peak is listed in parentheses. EC, extinction coefficient. QY, quantum yield. ND, not determined.

**Figure 2:**
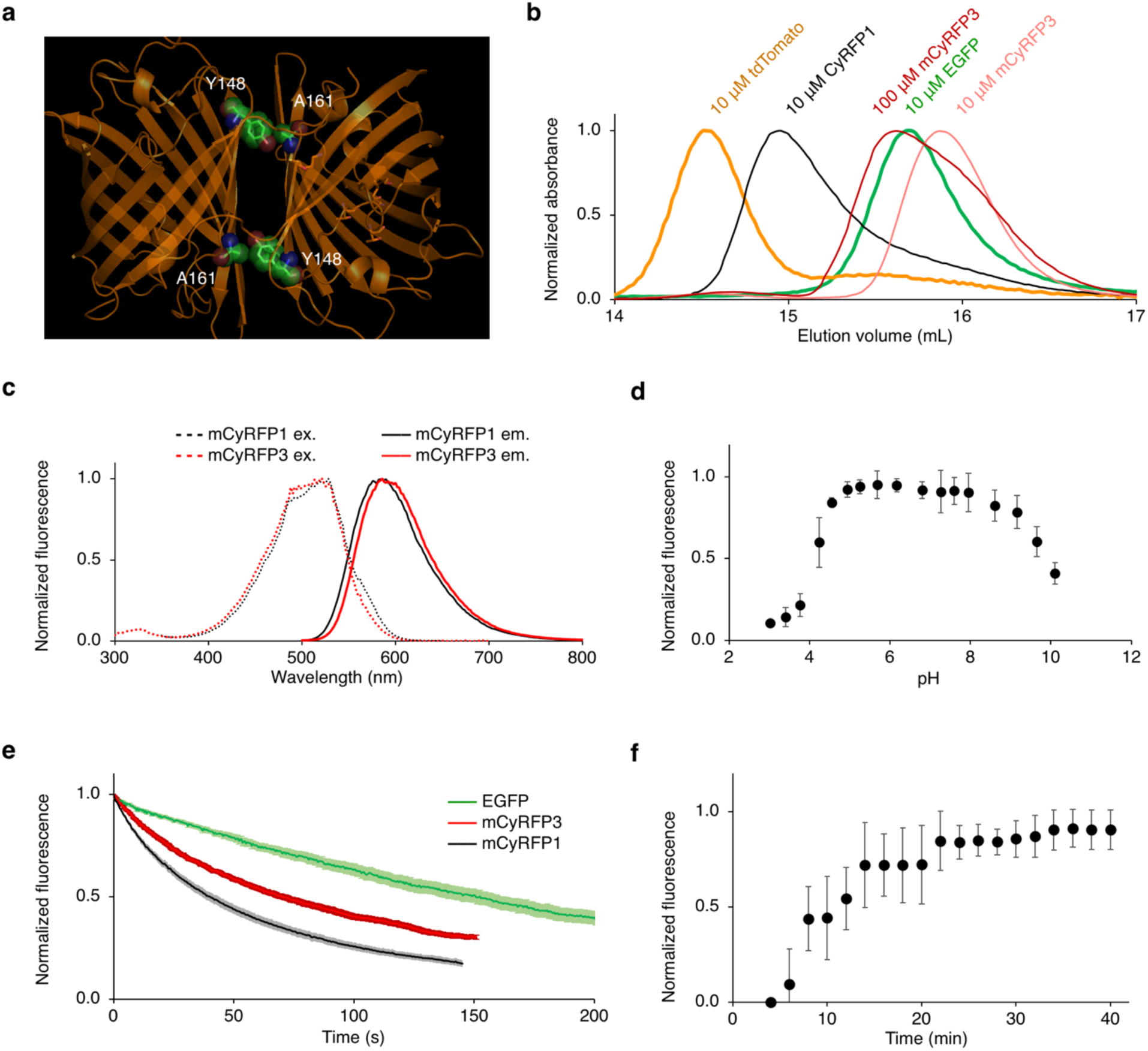
Engineering and characterization of mCyRFP3. Engineering of mCyRFP3. (a) Crystal structure of dimeric predecessor CyOFP1 displaying interacting residues Y148 and A161. (b) Size-exclusion chromatography of EGFP, tdTomato, CyRFP1, and mCyRFP3. EGFP was used as a monomeric standard, while tdTomato was used as a dimeric standard. (c) Excitation and emission spectra of mCyRFP3 compared to its parent mCyRFP1. (d) pH dependence of mCyRFP3 fluorescence, demonstrating a pKa of 4.1. Error bars are s.em.m of triplicate measurements. (e) Photobleaching kinetics of purified cyan-excitable red fluorescent proteins under illumination by a 120-W metal-halide arc lamp through a 490/20-nm excitation filter. The time-axis was adjusted for each fluorophore to simulate excitation conditions producing 1000 photons per s per molecule. Lighter shading on CyRFP1, mCyRFP1, and mCyRFP3 lines represents standard deviation of five measurements. (f) Maturation kinetics of mCyRFP3 demonstrating a half-life of t = 12.5 min.

We next characterized mCyRFP3 performance in mammalian cells. Fusions to various subcellular proteins were properly localized (**Fig. 3a)** and a histone H2B fusion enabled observation of mitosis **(Fig. 3b)**. In the organized smooth endoplasmic reticulum (OSER) assay, mCyRFP3 showed a similar score (87% whorl-free cells) as mCyRFP1 **(Fig. 3c)**. While purified proteins *in vitro* provide extinction coefficients and quantum yield parameters as an objective measure of a mature FP’s per-molecule brightness, an FP’s effective brightness in cells is influenced by its maturation efficiency and stability. We compared the brightness of cells expressing mCyRFP3, mCyRFP1, or CyRFP1 using a bicistronic construct co-expressing EGFP, where the EGFP fluorescence was used to normalize for differences in transfection and mRNA levels. In HEK293A and HeLa cells, mCyRFP3 was significantly brighter than mCyRFP1 **(Fig. 3d,e)**.

**Figure 3:**
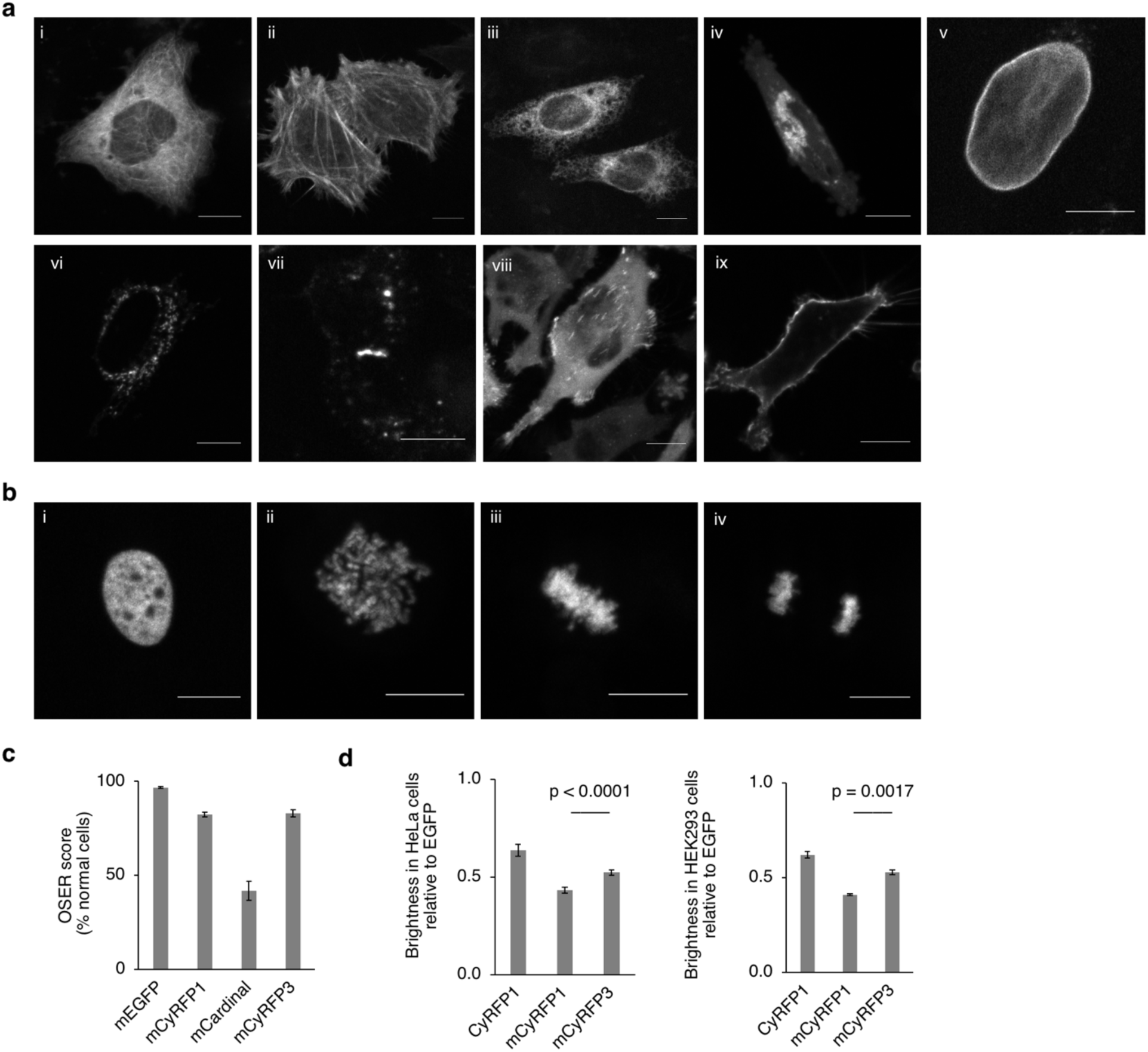
Performance of mCyRFP3 in mammalian cells. (a) HeLa cells expressing mCyRFP3 fused various subcellular proteins. For each fusion, the original of the fusion partner and its normal subcellular location are indicated in parentheses. (i) mCyRFP3-2aa-tubulin (human, microtubules), (ii) mCyRFP3-7aa-actin (human, actin cytoskeleton), (iii) Calnexin-14aa-mCyRFP3 (human, endoplasmic reticulum), (iv) mannosidaseII-10aa-mCyRFP3 (mouse, Golgi complex), (v) mCyRFP3-10aa-lamin B1 (human, nuclear envelope) (vi) PDHA-10aa-mCyRFP3 (human, mitochondrial pyruvate dehydrogenase), (vii) connexin43-7aa-mCyRFP3 (rat, cell-cell adhesion junctions), (viii) paxillin-22aa-mCyRFP3 (chicken, focal adhesions), (ix). mCyRFP3-2aa-CAAX. Scale bar, 10 µm. (b) mCyRFP3-10aa-H2B (human, nucleosomes) in (i) interphase, (ii) prophase, (iii) metaphase, (iv) anaphase. (c) Performance of mCyRFP3 compared to mCyRFP1, mEGFP and mCardinal on CytERM monomericity assay. Error bars represent standard deviation of measurements from > 150 cells in each of three separate experiments. (d) mCyRFP3 characterization in mammalian cells. Brightness comparison of mCyRFP3 in HEK293A and HeLa cells expressing a bicistronic construct EGFP-P2A-RFP where RFP = CyRFP1, mCyRFP1, and mCyRFP3. The red fluorescence generated from each of the RFPs were normalized to the fluorescence of EGFP to normalize for expression and were charted relative to the value of CyRFP1. Excitation was performed with a 480/10-nm filter and emission was collected from 580 to 800 nm. Integrated red emission relative to the integrated GFP emission is shown as mean ± s.e.m. of 6–8 biological replicates.

To determine if, optimization of amino acids 148 and 161 could also improve the performance of another related RFP, we also performed combinatorial saturation mutagenesis at these positions in the mMaroon1 far-red fluorescent protein^13^. Indeed simultaneous Y148T A161G mutations led to the development of a more monomeric and brighter variant, mMaroon2 **(Supplementary Note, Supplementary Figs 1–2, Supplementary Table 1)**.

We recently developed ASAP3, a GEVI with improved responsivity and brightness compared to previous ASAP-family GEVIs^5^. An independent comparison found that ASAP3 exhibits the largest power-normalized signal-to-noise ratios for both subthreshold voltage dynamics and action potentials (APs) among GEVIs developed to date^14^. We thus explored whether mCyRFP3 could serve as a non-perturbing voltage-independent label for ASAP3. We tested fusions of ASAP3 to mCyRFP3, mCyRFP1, mRuby3, which we had previously tested with ASAP1, and the recently developed mScarlet, which is non-perturbing in fusions with a variety of cellular proteins^15^. ASAP3-mCyRFP3 (ASAP3-R3) showed the clearest membrane localization in dendrites and the smallest amount of cell body accumulations, followed by ASAP3-mCyRFP1, then ASAP3-mRuby3, and finally ASAP3-mScarlet, which completely lacked membrane expression and whose cell body accumulations were especially large (**Fig. 4a)**. High-magnification confocal microscopy confirmed tight correspondence between the green signal of ASAP3 and the red signal of mCyRFP3 (**Fig. 4b**). Conveniently, both fluorophores of ASAP3-R3 can be excited by the same wavelength, while their emissions are easily separable (**Fig. 4c)**. The steady-state voltage responsivity of ASAP3-R3 in the green channel upon depolarization from –70 mV to 30 mV was –41 ± 1% (mean ± standard error of the mean, n = 8 HEK293 cells) (**Fig. 4d**). The slightly lower response amplitude compared to ASAP3 (–51 ± 1%)^5^ may be due to minor contamination by voltage-insensitive CyRFP2 fluorescence in the green emission channel (**Fig. 4c**).

**Figure 4:**
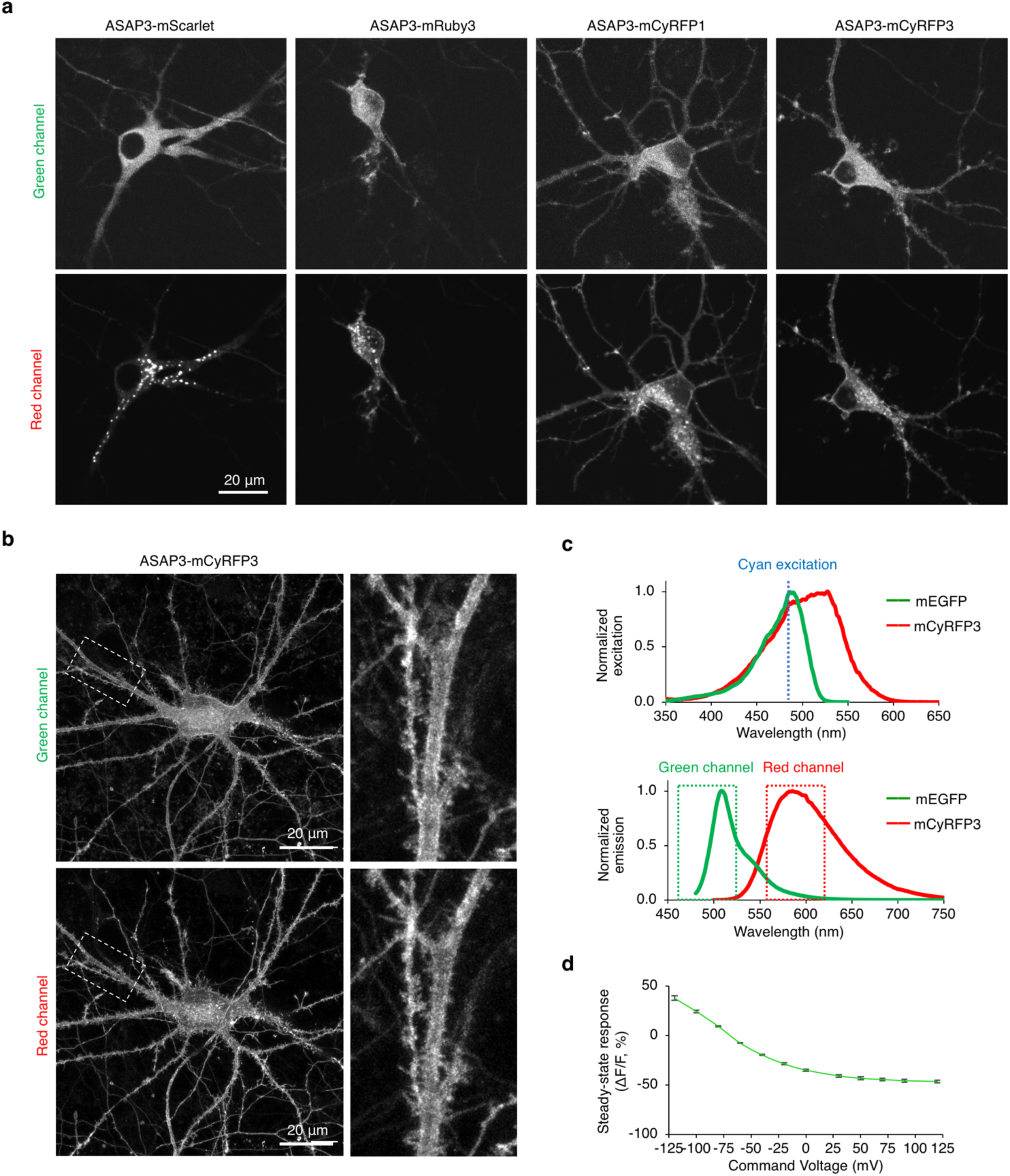
ASAP3-R3. (a) Representative epifluorescence images of ASAP3 fusions to various RFPs expressed in primary rat hippocampal neurons. (b) Confocal images of ASAP3-R3 in a rat hippocampal neuron. (c) Above, normalized excitation spectra of mCyRFP3 and mEGFP shows single cyan excitation wavelength can be used. Below, normalized emission spectra of mCyRFP3 and mEGFP demonstrates easily separable fluorescence. (d) Steady-state voltage responsivity of ASAP3-R3. Error bars represent standard error of the mean (n = 8 HEK293 cells).

Cardiomyocytes (CMs) derived from induced pluripotent stem cells (iPSCs) enable the study of patient-specific mechanisms in cardiovascular diseases^16^. A key functional readout of iPSC-CMs is the pattern of spontaneous cardiac APs, which drive contraction and whose appearance indicates the maturation of ion homeostasis and channel properties. Recording APs is also useful for studying signaling pathways leading to lineage specification of CM subtypes^17^. The gold standard for characterizing APs in iPSC-CMs is patch-clamp electrophysiology, but this method is low-throughput and destructive. An alternative method for AP measurements is to use fluorescent GEVIs, but GEVIs preferentially would be ratiometric so that changes in membrane potential are self-normalized and can be discriminated from motion-induced changes in fluorescence.

We tested the utility of ASAP3-R3 in reporting APs of iPSC-CMs. ASAP3-R3 fluorescence from beating iPSC-CMs was excited using blue light, then green and red channels were recorded by a single CMOS camera using an emission beam splitter. We found some apparently non-motile iPSC-CMs were firing APs, as ASAP3-R3 showed rhythmic decreases in fluorescence in the green channel and no change in the red channel **(Fig. 5a)**. ASAP3-R3 showed a response amplitude of ΔR/R of –46.6 ± 4.8% compared with –47.8 ± 5.21% for ASAP3 in spontaneously beating iPSC-CMs **(Fig. 5b)**, demonstrating the ASAP3-R3 GEVI is fully functional. Photobleaching was not observed in the recording period (**Fig. 5b**).

**Figure 5:**
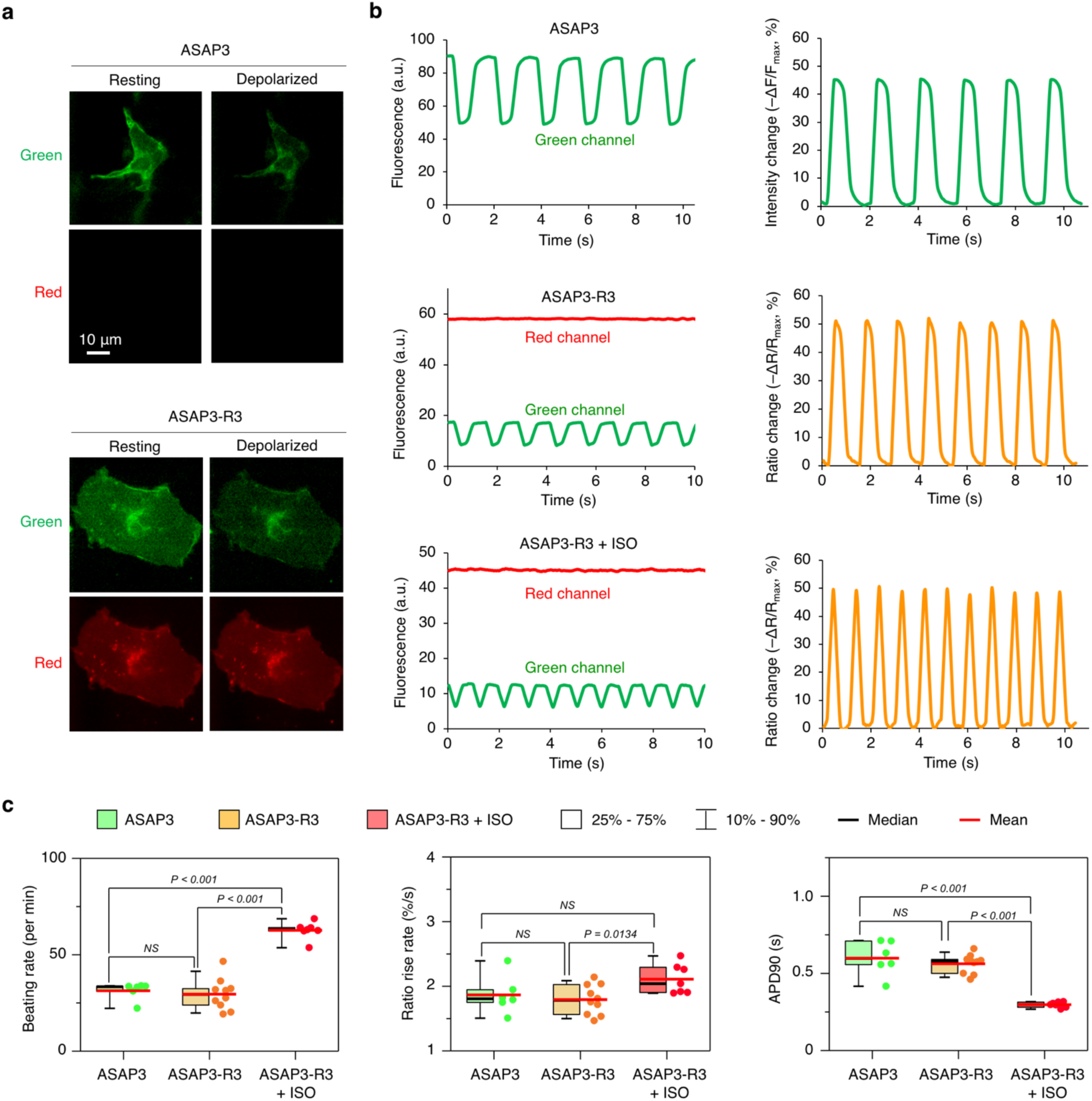
ASAP3-R3 performance in cardiomyocytes. (a) Representative induced pluripotent stem cell derived cardiomyocytes (iPSC-CMs) expressing ASAP3 of ASAP3-mCyRFP. (b) Representative single-trial ΔR/R or ΔF/F responses to spontaneous cardiac APs of ASAP3, ASAP3-R3, cells expressing ASAP3-R3 treated with isoproterenol (ISO). (c) Characterization of traces obtained from cells expressing ASAP3 (green, n = 6), ASAP3-R3 (orange, n = 10), and ASAP3-R3 + ISO (salmon, n = 7).

We also tested if ASAP3-R3 can faithfully report drug induced changes of APs in iPSC-CMs. Isoproterenol (ISO) activates β-adrenergic receptor in CMs to increase heart rate. ASAP3-R3 showed that ISO treatment of iPSC-CMs also increased the spontaneous contraction rate, from ∼30 beats per minute (bpm) to > 60 bpm (**Fig. 5b,c**). The rise rate, which should correlate with the up-stroke velocity in patch clamp measurement, was also significantly increased (**Fig. 5c)**. As a result, the AP duration to 90% repolarization (APD90) was significantly shorter in the ISO treated group (**Fig. 5c)**.

We next demonstrated how ASAP3-R3 can be used to correct for motion artifacts in moving cells. Different edges of a contracting iPSC-CM can exhibited disparate responses in green intensity **(Fig. 6a,b)**. On one edge, green signal fluctuations were larger than those in non-contracting CMs (**Fig. 6a**). On another edge, the green signal was inverted from the expected response and did not resemble AP waveforms (**Fig. 6b)**. The voltage-insensitive red channel showed different rhythmic fluctuations at the two edges, indicating different patterns of membrane movement. Thus, the disparate green fluorescence responses from the same iPSC-CM were due to different contributions of membrane movement to green fluorescence. Indeed, when the green/red ratio was tracked instead, different portions of the membrane responded in the same direction and magnitude as in non-contracting iPSC-CMs (**Fig. 6a,b**). The ratio changes in different membrane regions also resembled each other in waveform shape even though the individual channels did not (**Fig. 6a,b**).

**Figure 6:**
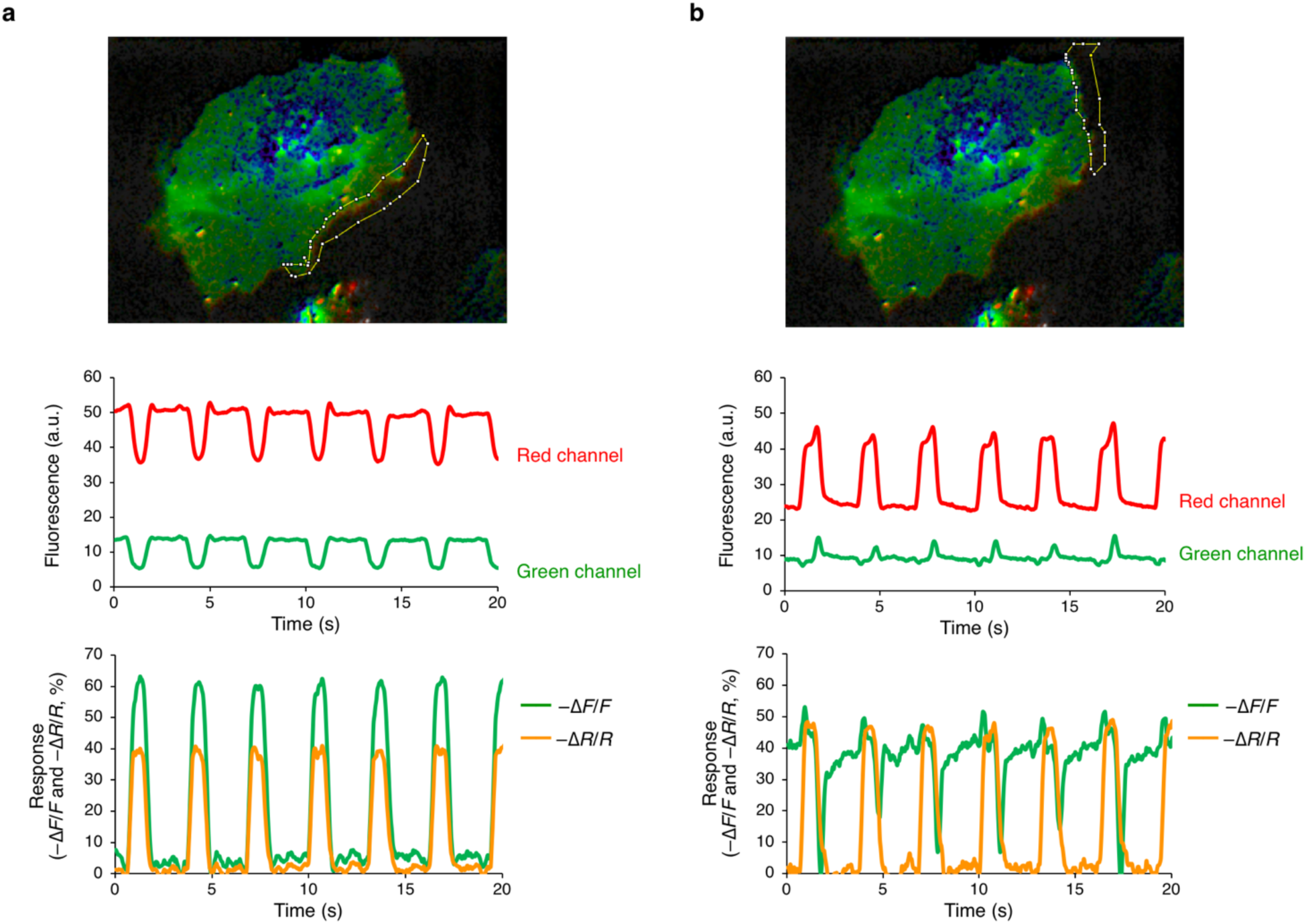
The red channel in ASAP3-R3 allows for motion correction. (a) Top, one segment of membrane in a beating CM was analyzed. Middle, raw traces of green and red signals in the selected region. Red trace indicates significant cell movement in and out of selected region detected by the voltage-independent mCyRFP. Bottom, relative change in green fluorescence alone (–ΔF/F) and the relative change in the green/red ratio (–ΔR/R). While green intensity changes are larger than those observed previously in non-contracting CMs due to these motion artifacts, ratio changes are similar in magnitude to those observed prevoiusly. (b) Similar analysis on a different membrane segment of the same cell. Here, the green fluorescence changes are opposite in direction from that expected due to movement, which is detected in the red channel. While green intensity traces do not resemble AP waveforms, ratio traces are indistinguishable from AP waveforms in non-motile CMs and similar in shape to those in (a).

Finally, using ASAP3-R3, we demonstrated regulation of the membrane potential during the cell cycle. Potassium and chloride channel activities are regulated by cell cycle phase^18^, and electrode measurements have found fluctations in the transmembrane potential during cell cycle progression^19,20^. Interestingly, manipulations of channel function or extracellular ion concentrations suggest a role for transmembrane voltage in the timing of cell cycle transitions^18^. However, direct electrical measurements across the cell cycle have only been reported as single-timepoint measurements in non-motile cell types. We asked whether we could observe transmembrane voltage in a motile cell type throughout an entire cell cycle. HeLa cells demonstrate pronounced cellular motility, which would make repeated membrane voltage measurements of a single cell throughout the cell cycle difficult. The morphological dynamism of HeLa cells would also prevent accurate voltage readings from a non-ratiometric GEVI.

We thus expressed ASAP3-R3 in HeLa cells and visualized the green/red ratio over time. We observed a ∼35% change in green/red ratios throughout the cell cycle, with the lowest ratio value (representing depolarization) in mitosis prior to cytokinesis (**Fig. 7**). The highest green/red ratios (representing hyperpolarization) occurs 6–9 hours following cytokinesis, which corresponds to S-phase in HeLa cells^13^. The timing of these voltage changes are consistent with previous reports using intracellular electrodes on non-motile cell types^19,20^. Thus, using ASAP3-R3, we were able to detect membrane voltage changes throughout the cell cycle in moving cells, finding that HeLa cells are most depolarized during mitosis and most polarized during S-phase.

**Figure 7:**
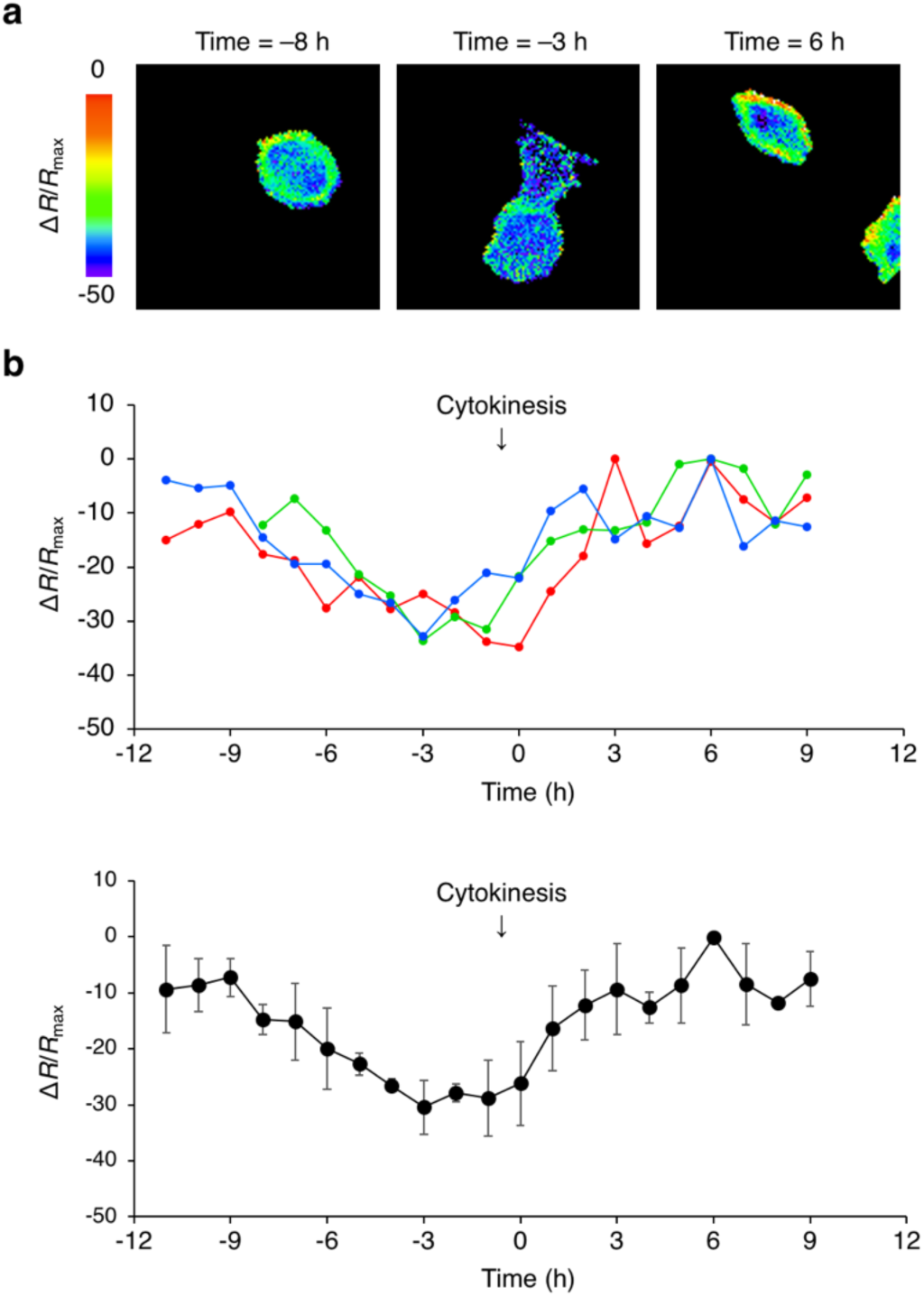
Observing membrane potential changes during the cell cycle with ASAP3-R3. (a) Images of the ASAP3-R3 green/red ratio at different time points relative to cytokinesis in a HeLa cell. (b) Above, individual green/red ratio time courses of three HeLa cells during cell division. The cell in (a) is represented by the blue trace. Cytokinesis occurs immediately before the 0-h time point. Below, mean green/red ratios of the three cells. Error bars represent standard deviation.

## Discussion

A high-performance ratiometric GEVI would be highly desirable to allow transmembrane voltage imaging in specimens under motion, as changes in location or focus would alter the intensity of a single channel but not the ratio between channels. After various RFPs fused to ASAP-family GEVIs interfered with membrane expression, we developed mCyRFP3 as a brighter and more monomeric cyan-excitable red fluorescent protein from hypothesis-driven combinatorial saturation mutagenesis. ASAP3-R3 proved to be well expressed and to produce voltage-modulated ratios that are resistant to motion artifacts. We demonstrated ASAP3-R3 can report AP waveforms even at motile regions of contracting cardiomyocytes. We also used ASAP3-R3 to track cyclical fluctuations in membrane voltage during the cell cycle of HeLa cells, discovering depolarization during mitosis and hyperpolarization during S-phase.

As mCyRFP3 is more monomeric *in vitro* than mCyRFP1, it is not surprising that it would be less perturbing to ASAP3 membrane localization than mCyRFP1. However, in the OSER assay, fusions of mCyRFP1 and mCyRFP3 to CytERM showed similar propensities to form ER whorls. In addition, although FusionRed and mScarlet are less perturbing in the OSER assay than mCyRFP3, they performed worse when fused to ASAP3. These observations demonstrate that the OSER assay only tests the performance of fluorescent proteins fused to one cellular protein, and may not necessarily predict performance of fusions with any other particular protein. One possible complicating factor in the OSER assay is protein expression level; fusions with less stable fluorescent proteins may accumulate to lower concentrations and thereby create whorls in a smaller percentage of cells.

Our results demonstrate the utility of structure-guided multi-site saturation mutagenesis for fluorescent protein improvement. Obtaining mCyRFP3 would have been exceedingly difficult by random mutagenesis. mCyRFP3 differs by mCyRFP1 by three nucleotides within a 700 bp gene, so in a library bearing 3 random mutations per clone without duplicates, this combination would have occurred once in 700 choose 3 × 4^3^, or 3.6 × 10^9^ mutants. In reality, the library would need to be at least an order of magnitude larger to account for Poisson distributions of mutation combinations. If the ideal combination of amino acids had required six nucleotide changes, then it would have arisen in a perfectly distributed library of six-nucleotide mutants once per 6.6 × 10^17^ clones. By identifying two sites on the crystal structure, we only needed to screen for 1024 possible combinations using an NNS codon at each site (32 × 32). Thus, hypothesis-driven structure guided mutagenesis can identify beneficial multi-site mutants that would have been unobtainable using random mutagenesis. Similar methods have been used to produce functional improvements in a variety of RFPs and in ASAP3 itself^5,7,10,13,21,22^.

Indeed, our observation that coordinated mutations at positions 148 and 161 produced more monomeric and brighter variants of multiple RFPs suggests a general strategy of monomerization in which positions interacting across a dimer interface are mutated to all possible combinations. While this may seem obvious in retrospect, this does not appear to be a standard step in the monomerization workflow for fluorescent proteins. mCyRFP1 is derived from TagRFP, which in turn is a weakly dimeric mutant of the natural tetrameric RFP eqFP578. To our knowledge, positions 148 and 161, which are distant from each other on the surface of a single monomer, have not been previously mutated in combination and to saturation in any RFPs derived from TagRFP.

In summary, we have created a cyan-excitable red fluorescent protein, mCyRFP3, with improved brightness and monomericity. Among RFPs, mCyRFP3 performs especially well as a fusion tag for the ASAP3 voltage indicator. The resulting ASAP3-R3 enables voltage imaging with an internal reference channel for motion correction, as we demonstrate by imaging transmembrane voltage changes during cardiomyocyte contraction and during the cell cycle. Compared to voltage indicator dyes, ASAP3-R3 has the advantage of being genetically encoded, allowing for stable expression and targeting to specific cell types. We expect that ASAP3-R3 will also be useful for imaging voltage in the brains of living animals, where the mCyRFP3 reference channel can allow for correction of motion artifacts in the voltage-modulated ASAP3 signal.

## ACKNOWLEDGEMENTS

We thank Lin Ning (Lin laboratory) for assistance with organotypic slice cultures and for Jonathan Mulholland (Stanford Cell Sciences Imaging Facility) for assistance with confocal microscopy. This work was supported by a NSF Graduate Research Fellowship (B.B.K.), the Stanford Department of Bioengineering (Y.A.H., B.B.K., M.C.), grant NNF18OC0031816 from the Novo-Nordisk Foundation and the Stanford Bio-X Program (M.W.), the Vitalant and Hemophilia Center of Western Pennsylvania (H.W.), NIH grants K99 HL133473 (H. W.), R01HL126527 (J.C.W.), 1R01HL133272 (J.C.W. and M.Z.L.), 1U01NS090600 (M.Z.L.), 1U01NS103464 (M.Z.L.), and 1RF1MH114105 (M.Z.L.).

## AUTHOR CONTRIBUTIONS

B.B.K. cloned and characterized mCyRFP3 and ASAP3 fusions, imaged and analyzed cardiomyocyte experiments, prepared figures, and wrote the manuscript. Y.A.H. designed, performed, and analyzed cell cycle experiments. M.P. cloned and characterized ASAP1 fusions. M.C. conceived of experiments and cloned and characterized ASAP1 fusions. H.W. generated cardiomyocytes and analyzed cardiomyocyte imaging experiments. M.W. assisted with cardiomyocyte imaging and analysis. F.S.-P. supervised the design and analysis of ASAP1 fusions. J.C.W. supervised cardiomyocyte generation. M.Z.L. supervised all experiments, assisted with data analysis, prepared figures, and wrote the paper.

## COMPETING FINANCIAL INTERESTS

The authors declare no competing financial interests.

## METHODS

### Mutagenesis and screening of libraries

Plasmids were constructed using standard molecular biology methods including polymerase chain reaction (PCR) and In-fusion cloning (Clontech). Mutations for specific residues were introduced by overlap-extension PCR. All cloning junctions and PCR products were sequence verified. Mutants were expressed and screened in constitutively active bacterial expression vector pNCS (Allele Biotech). Plasmids were transformed into chemically competent XL-10 Gold (Agilent), and colonies were grown on LB agar plates at 37 °C for 16-20 h and at room temperature for an additional 20-24 h. For each round of mutagenesis, the number of colonies screened was 10-fold the expected library diversity to ensure full coverage. Colonies expressing mCyRFP3 variants were screened for transmitted color by eye and for fluorescence in a Fluorochem Q imaging enclosure (Alpha Innotech) with an 475/42-nm excitation filter and an 699/62-nm emission filter.

### Protein production and characterization

For spectral characterization, bacterial pellets were lysed in B-PER II (Pierce) and hexahistidine-tagged proteins were purified with HisPur Cobalt Resin (Pierce). Proteins were desalted into phosphate-buffered saline (PBS) pH 7.4 using Econo-Pac 10DG gravity flow chromatography columns (Bio-Rad). Absorbance, excitation spectra, and emission spectra were measured with Safire2 or Infinite M1000 Pro plate readers (TECAN). Extinction coefficients were calculated using the base-denaturation method^21^. Quantum yields for mCyRFP3 were determined in PBS at pH 7.4 by exciting with 475- to 485-nm light and integrating emission from 500 to 800 nm, corrected for detector sensitivity. mCyRFP1 was used as the quantum yield standard. *In vitro* photobleaching measurements were performed in PBS droplets under mineral oil on an IX81 inverted microscope with a 40× 1.15-numerical aperture (NA) water-immersion objective, an X-Cite 120-W metal halide lamp (Lumen Dynamics) at 100% neutral density, a 485/10-nm excitation filter (Omega), and an Orca ER camera (Hamamatsu) controlled by Micro-Manager software^23^. Images were acquired every 1 s under continuous illumination. Times were scaled to produce photon output rates of 1000 per molecule per s as previously described^24^. pH titration was performed using a series of buffers (1M HOAc, 1M NaOAc, 5M NaCl for pH 3–5.5; 100 mM KH_2_PO_4,_ 100 mM K_2_HPO_4_ for pH 6-8; 100 mM glycine for pH 9.5-10). HCl or NaOH were used to adjust the pH. 5 μL of purified protein was diluted in 145 μL buffer with different pH values, and the fluorescence brightness was measured. Size exclusion chromatography was performed with a Superdex 200 30/100 GL column (GE Healthcare). 100 μL of 10–100 μM purified proteins were loaded and eluted at a flow rate of 0.5 mL/min. Protein elution was monitored by absorbance at 280 nm.

### Brightness comparison of fluorescent proteins in mammalian cells

HeLa (ATCC) and HEK293A (GE Dharmacon, Fischer Scientific) cells were grown on glass-bottom dishes (Cellvis) in high-glucose Dulbecco’s Modified Eagle Medium (DMEM, Hyclone) supplemented with 10% fetal bovine serum (Gemini), 2 mM glutamine (Gemini), 100 U/mL penicillin and 100 μg/mL streptomycin (Gemini) to 70-80% confluency, then transfected using Lipofectamine 2000 (Thermo Fisher) with a EGFP-P2A-RFP-CAAX construct, where RFP was CyRFP1, mCyRFP1, or mCyRFP3. EGFP served to normalize for transfection efficiency and cell number. One day post-transfection, approximately 10^5^ cells were replated in a 96-well plate and emission spectra from 500 to 800 nm, upon excitation at 480 nm with a bandwidth of 10 nm, were obtained in an Infinite M1000 Pro microplate reader (Tecan). As EGFP emission is negligible at peak RFP emission and vice versa, the EGFP-normalized RFP brightness was simply calculated as the ratio of RFP and EGFP peak values.

### Microscopy of fusion proteins

mCyRFP3 fusions were cloned into pLL3.7m, a modified form of pLL3.7, using standard molecular biology methods. Fusions were made to human calnexin (NM_001746.3), Lifeact, mouse mannosidase II (NM_008549.2), human laminB1 (NM_005573.2), human pyruvate dehydrogenase (NM_000284), chicken paxillin (NM_204984.1), human histone H2B (NM_021058.3), rat connexin-43 (NM_012326.2), and human α-tubulin (NM_006082). All sequences were gifts of M. Davidson (Florida State University). mCyRFP3 fused to the CAAX membrane-localization signals were subcloned into pcDNA3. HeLa cells were grown and transfected as above, then imaged 24–48 h later in FluoroBrite DMEM with B-27. mCyRFP3 fusions were imaged on an Axio Observer microscope with a 63 × 1.3-NA oil-immersion objective (Zeiss) equipped with an UltraVIEW spinning-disc confocal unit (Perkin-Elmer). Excitation was provided by a 488-nm laser excitation, and emission was collected through a 615/70-nm filter with an C9100-50 EMCCD camera (Hamamatsu). Maximal intensity projections of optical sections were generated in the ImageJ program ^25^.

### Organized smooth endoplasmic reticulum (OSER) assay

mCyRFP3, EGFP, and mCardinal were fused to the C-terminus of the signal-anchor transmembrane domain of cytochrome P450 (amino acids 1-29, CytERM) in a pcDNA3 vector using standard molecular biology methods. HeLa cells were grown and transfected as above, then imaged 18 h post-transfection in FluoroBrite DMEM with B-27 on the Axio Observer microscope with a 63× 1.3-NA objective (Zeiss) equipped with an UltraView spinning-disc confocal unit (Perkin-Elmer). The percentage of cells with visible whorls were counted, excluding cells with unusually bright expression. At least 150 cells were counted per dish, and three technical replicates using three dishes were performed. CytERM-EGFP and CytERM-mCardinal served as references for comparison to previously reported data^26^.

### Curve fittings for ASAP3-R3

HEK293-Kir2.1 cell lines^27^ were confirmed to have a polarized resting membrane potential of approximately –75 mV, and were grown on glass-bottom dishes (Cellvis) in high-glucose Dulbecco’s Modified Eagle Medium (DMEM, Hyclone), supplemented with 10% fetal bovine serum (Gemini), 2 mM glutamine (Gemini), and 400 µg/mL geneticin (Thermo Fisher). Cells were grown to ∼70-80% confluency and were transfected with ASAP3-R3 and different permutations of ASAP3, ASAP3-Y66A, and mCyRFP3 fusions with Kv2.1, using Lipofectamine 3000 (∼100 ng of DNA, 0.4 µL P3000 reagent, 0.4 µL Lipofectamine) followed by a change in media four hours later. Two days post-transfection, approximately 10^5^ cells were replated in a 96-well plate and emission spectra from 500 to 800 nm, upon excitation at 480 nm with a bandwidth of 10 nm were obtained on an Infinite M1000 Pro microplate reader. The spectra of the individual constructs containing either ASAP3 and mCyRFP3 were normalized to their peaks and were fitted using the sum of least squares to the combined spectra of the original fusion construct.

### Neuronal cell culture, transfection, and imaging

All procedures were approved and carried out in compliance with the Stanford University Administrative Panel on Laboratory Animal Care. Hippocampal neurons were extracted from embryonic day 18 Sprague Dawley rat embryos by dissociation in HBSS supplemented with 10 mM D-glucose and 10 mM HEPES pH 7.2 (Thermo Life Sciences). Digestion was performed in RPMI media containing 20 U/mL papain (Worthington Biochemical LS003119) and 0.005% DNAase I, at 37 °C for 15 min. Dissociated neurons were then plated in Neurobasal with 10% FBS, 2 mM GlutaMAX, and 2% B27 (Thermo Fisher) at a density of 4 × 10^6^ cells/cm^2^ in a 12-well glass-bottom plate, pre-coated overnight with 0.1 mg/mL > 300-kDa poly-D-lysine hydrobromide (Sigma). 6–8 h later media was replaced with Neurobasal with 1% FBS, 2 mM GlutaMAX, and B27, with refreshing media every 3– 4 days. Neurons were then transfected at 9–11 days *in vitro* with Lipofectamine 2000 (Thermo Fisher). 500 ng of total DNA (100 ng ASAP3-RFP and 400 ng pcDNA3) and 1.5 µL of Lipofectamine 2000 transfection reagent was used for each well. The culture medium from each well was first collected and stored at 37 °C and 5% CO_2_ and temporarily replaced with Neurobasal with 2 mM GlutaMAX. Neurons were imaged 3 days post-transfection on an inverted LSM 880 confocal microscope with a 40× 1.3-NA oil-immersion objective and Zen Blue software (Zeiss). Excitation was provided by a 488-nm laser, and emission was collected simultaneously at 493–566 nm and 565–656 nm. Maximal intensity projections of optical sections were generated in ImageJ.

### Differentiation, culture, transfection, and voltage imaging of human iPSCs derived cardiomyocytes

Stem cell protocols were approved from the Stanford University Human Subjects Research Institution Review Board. Human iPSC lines were maintained in 5% CO_2_ environment in Essential 8 Medium (Thermo Fisher). For cardiomyocyte differentiation, iPSCs were switched to insulin-free RPMI + B27, where they were treated with 6 µM CHIR99021 (Selleckchem) for 2 days, recovered for 1 day, and 5 µM IWR-1 (Sigma) for 2 days, recovered for another 2 days, and finally switched to RPMI+B27 plus insulin medium. After beating CMs were observed around day 9-11 of differentiation, iPSC-CMs were reseeded and purified with glucose-free RPMI+B27 medium for 2 rounds. The purity and quality of the iPSC-CMs were verified by FACS with TNNT2 and immune-staining of TNNT2 and α-actinin. The iPSC-CMs were maintained in RPMI + B27 medium until day 30 and were dissociated with TrypLE Select Enzyme (Thermo Fisher) and reseeded in glass-bottom dishes at a concentration of 50K cell per square centimeter. Briefly, 1 µg of plasmid ASAP3 and ASAP3-R3 were added into the solution of GeneJammer (Agilent) transfection reagent in Opti-MEM (3 µL in 200 µL), and incubated for 30 mins at RT. Then the mixtures were added directly into the cell culture and incubated for 12 hours. The cell cultures were allowed to incubate for an additional 3-4 days for optimal transfection results. Cells were then imaged at 20 Hz on an Axiovert 200M inverted microscope (Zeiss) with a 40× 1.2-NA C-Apochromat water-immersion objective and equipped with a stage-top incubator (LCI). Excitation from a 150-W xenon arc lamp (Zeiss) at 100% neutral density was passed through a 480/30 nm excitation filter (Chroma). Emissions were simultaneously collected in green (510/60 nm) and red (630/50) channels using a DualView image splitter (Mag Biosystems). Images were acquired on an ORCA-Flash4.0 V2 C11440-22CU scientific CMOS camera with a HCImage software (Hamamatsu). At least 10 cells were selected for recording. Fluorescence intensity in cellular regions of interest or a background region were measured in ImageJ. Mean background intensity was subtracted from cellular regions of interest.

## Supplementary Note

We hypothesized that combinatorial mutagenesis of Tyr-148 and Ala-161 could also improve monomericity and maturation of other TagRFP derivatives. The TagRFP derivative mMaroon1 features far-red emission that provides an additional channel distinct from orange fluorescent proteins^13^. However, mMaroon1 is only weakly monomeric, migrating on native polyacrylamide gel electrophoresis as a monomer but eluting as a dimer in size-exclusion chromatography (SEC) at 10 µM (**Supplementary Fig. 1a**). We thus performed combinatorial saturation mutagenesis of positions 148 and 161 in mMaroon1 to improve its monomericity and maturation. Indeed, we identified one mutant, mMaroon1 Y148T A161G, that was as bright as mMaroon1 in bacteria and exhibited improved monomericity, eluting in SEC as a dimer-monomer mixture at 10 µM (**Supplementary Fig. 1a**).

To further enhance monomericity, we explored removal of the 20-residue C-terminal tail, which engages in an dimerizing interaction in crystal structures of Neptune1, the most recent evolutionary predecessor to mMaroon1 for which a crystal structure exists^22^. In SEC, mMaroon1 ΔCT at 10 µM eluted in a broad peak overlapping with the dimeric standard (**Supplementary Fig. 1b**). Interestingly combining Y148T A161G mutations and tail deletion created a protein that was monomeric at 100 µM (**Supplementary Fig. 1b**). However, mMaroon1 Y148T A161G ΔCT was much dimmer in bacteria than mMaroon1 (**Supplementary Fig. 1c)**. This was despite similar chromophore maturation efficiency as a fraction of purified protein (75% for mMaroon1 Y148T A161G ΔCT vs. 79% for mMaroon1), suggesting protein instability rather than slow maturation as the cause of low fluorescence. Random mutagenesis of mMaroon1 Y148T A161G ΔCT identified an E175K mutant as maturing well in bacteria **(Supplementary Fig. 1c)** while maintaining monomericity at 100 µM **(Supplementary Fig. 1d)**. This protein, designated mMaroon2, exhibited peak absorbance and excitation at 604–606 nm (**Supplementary Fig. 1e,f**), a slightly red-shifted emission peak at 658 nm **(Supplementary Fig. 1f)**, and an extinction coefficient of 81 mM^-1^ cm^-1^ **(Table 1)**. Excitation by cyan light revealed a smaller residual green component compared to mMaroon1 **(Supplementary Fig. 1g)**. mMaroon2 displayed a p*K*_a_ of 6.1 **(Supplementary Fig. 1h)**, a maturation half-time of t = 8 min **(Supplementary Fig. 1i)**, and a normalized photobleaching half-time of 90 s **(Supplementary Fig. 1j)**.

We characterized the performance of mMaroon2 in mammalian cells. A fusion of histone 2B with mMaroon2 did not interfere with mitosis **(Supplementary Fig. 2a)** and fusions of various subcellular proteins to mMaroon2 were properly localized **(Supplementary Fig. 2b)**. We compared the brightness of cells expressing the parent dimer mMaroon1, mMaroon1 Y148T A161G ΔCT, and mMaroon2 from a bicistronic construct co-expressing mTurquoise2, using mTurqouise2 fluorescence to normalize for any differences in transfection efficiency or mRNA stability. In HEK293A cells, mMaroon2 was as bright as mMaroon1 and five-fold brighter than mMaroon1 Y148T A161G ΔCT **(Supplementary Fig. 2c)**. In HeLa cells, mMaroon2 was almost two-fold brighter than mMaroon1 and eight-fold brighter than mMaroon1 Y148T A161G ΔCT **(Supplementary Fig. 2c)**.

**Supplementary Table 1.** Engineering of mMaroon2.

|  | Peak<br>excitation<br>(nm) | Peak EC<br>(mM<sup>-1</sup>cm<sup>-1</sup>) | Peak<br>emission<br>(nm) | QY | Brightness<br>(M<sup>-1</sup>cm<sup>-1</sup>) | Multimericity<br>(at 10 μM) |
| --- | --- | --- | --- | --- | --- | --- |
| mMaroon1 | 608 | 80 | 654 | 0.11 | 8.8 | Dimer |
| mMaroon1 Y148T<br>A161G | 606 | 90 | 656 | 0.10 | 9.0 | Monomer-<br>dimer mixture |
| mMaroon1 ΔCT | 606 | 93 | 660 | 0.11 | 10 | Monomer-<br>dimer mixture |
| mMaroon1 Y148T<br>A161G ΔCT | 604 | 79 | 658 | 0.10 | 7.9 | Monomer |
| Maroon1 Y148T A161G<br>E175K ΔCT (mMaroon2) | 604 | 81 | 658 | 0.10 | 8.1 | Monomer |

EC, extinction coefficient. QY, quantum yield.

**Supplementary Figure 1.**
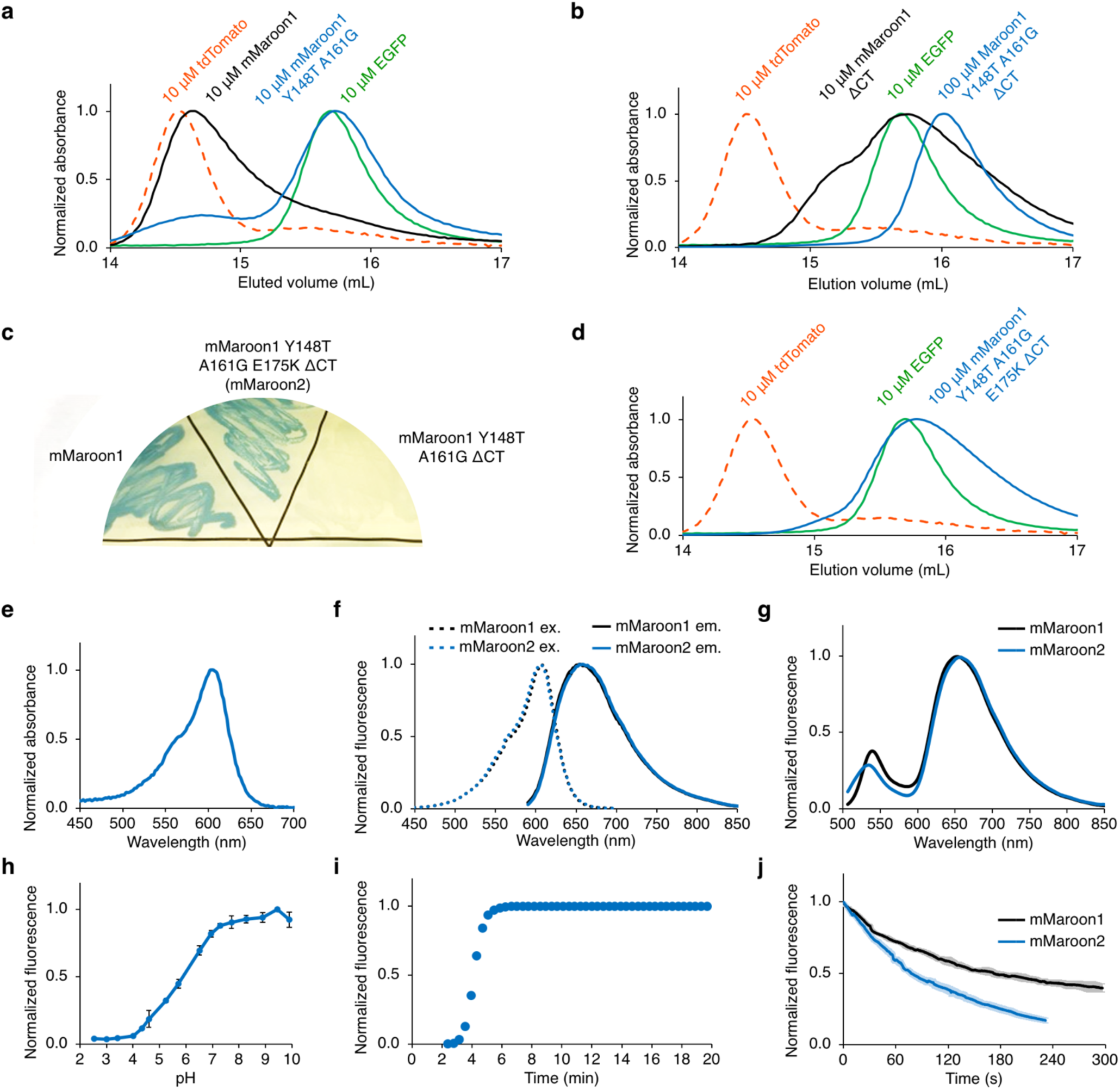
Characterization of mMaroon2. (a) In SEC, mMaroon1 elutes as a dimer while mMaroon1 Y148T A161G elutes as a dimer-monomer mixture at 10 μM concentration. EGFP was used as a monomeric standard, while tdTomato was used as a dimeric standard. (b) In SEC, 10 μM mMaroon1 ΔCT elutes with a broad size distribution while 100 μM mMaroon1 Y148T A161G ΔCT (mMaroon1.1) elutes as a monomer. (c) Patches of bacteria expressing mMaroon1, mMaroon1 T148 G161 K175 ΔCT (mMaroon2), and mMaroon1 T148 G161 ΔCT grown for 2 d. (d) In SEC, 100 μM mMaroon1 T148 G161 K175 ΔCT (mMaroon2) elutes as a monomer. (e) Absorbance of mMaroon2. (f) Excitation and emission of mMaroon2. (g) Emission of mMaroon1 and mMaroon2 excited at 480/10 nm and collected from 500 nm to 800 nm. (h) The pKa of mMaroon2 was measured as 6.1. Error bars are s.e.m of triplicate measurements. (i) Maturation kinetics of mMaroon2. (j) Photobleaching kinetics of purified mMaroon1 or mMaroon2 with a 120-W metal-halide arc lamp through a 615/30-nm excitation filter. The time axis was adjusted for each fluorophore to simulate excitation conditions producing 1000 photons per s per molecule. Lighter shading represents standard deviation of five measurements.

**Supplementary Figure 2:**
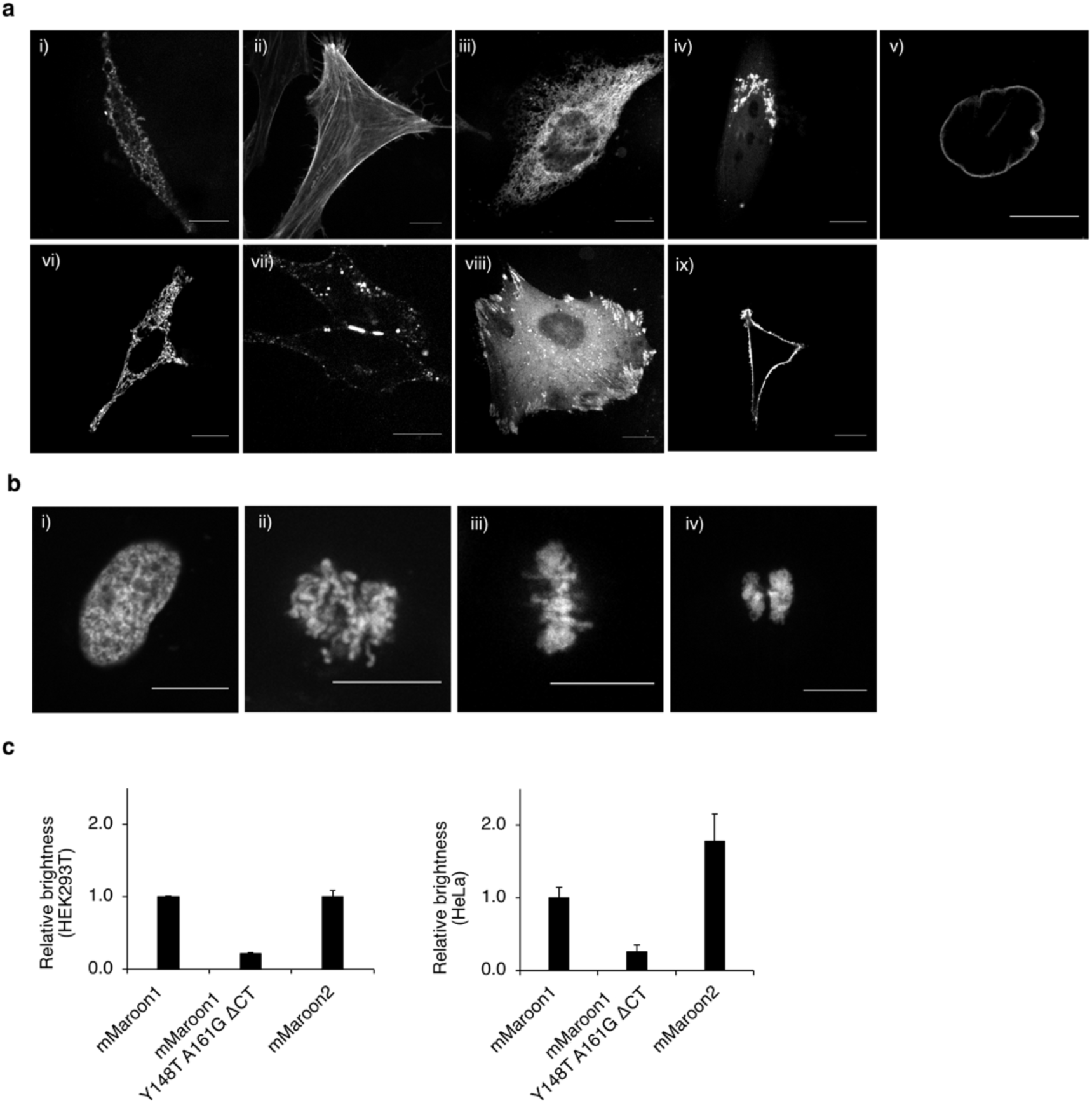
Performance of mMaroon2 Fusions. (a) HeLa cells expressing mMaroon2 fused to various domains. For each fusion, the original of the fusion partner and its normal subcellular location are indicated in parentheses. (i) mMaroon2-2aa-tubulin (human, microtubules), (ii) mMaroon2-7aa-actin (human, actin cytoskeleton), (iii) Calnexin-14aa-mMaroon2 (human, endoplasmic reticulum), (iv) mannosidaseII-10aa-mMaroon2 (mouse, Golgi complex), (v) mMaroon2-10aa-lamin B1 (human, nuclear envelope) (vi) PDHA-10aa-mMaroon2 (human, mitochondrial pyruvate dehydrogenase), (vii) connexin43-7aa-mMaroon2 (rat, cell-cell adhesion junctions), (viii) paxillin-22aa-mMaroon2 (chicken, focal adhesions), (ix). mMaroon2-2aa-CAAX. Scale bar, 10 µm. (b) mMaroon2-10aa-H2B (human, nucleosomes) in (i) interphase, (ii) prophase, (iii) metaphase, (iv) anaphase. (c) Brightness comparison of mMaroon2 in HEK293A and HeLa cells expressing the bicistronic construct mTurquoise2-P2A-RFP where RFP is mMaroon1, mMaroon1 Y148T A161G ΔCT, or mMaroon2. The red fluorescence generated from each of the RFPs was divided by the fluorescence of mTurquoise2 to correct for transfection and then normalized to the value of mMaroon1. Graphs are mean ± s.e.m. from six measurements.

## Notes

### Competing Interest Statement

The authors have declared no competing interest.

### Summary of Updates

mCyRFP2 name changed to mCyRFP3, as a different mCyRFP2 was recently published

